# Mapping Allosteric Communication in the Nucleosome with Conditional Activity

**DOI:** 10.1101/2025.09.14.676121

**Authors:** Augustine C. Onyema, Chukwuebuka Dikeocha, Jonathan Moussa, Sharon M. Loverde

## Abstract

The nucleosome core particle (NCP) regulates genome accessibility through dynamic allosteric communication between histone proteins and DNA. Building on the concept of conditional activity introduced by Lin (2016), we use molecular dynamics simulations and develop an open-source Python library, CONDACT (CONDitional ACTivity), to quantify time-resolved kinetic correlations in nucleosome systems. We analyze long-time simulations of the nucleosome core particle, including two different DNA sequences, the Widom-601 and ASP (alpha-satellite palindromic) sequences. By tracking dihedral angle transitions, we identify residues with high dynamical memory and map inter-residue communication pathways across histone subunits and DNA. Our analysis reveals kinetically connected domains involving post-translational modification sites, oncogenic mutation sites, and DNA contact regions, with dynamic coupling observed over distances up to 7.5 nm. These findings offer new insight into the long-range allosteric behavior of the nucleosome and its potential role in regulating chromatin accessibility. Quantifying this allosteric behavior potentially identifies targetable residues and domains for therapeutic intervention.

**Statement of Significance:** The nucleosome regulates DNA accessibility and plays a key role in epigenetic control of gene expression, yet the kinetic mechanism by which local modifications influence distant regions remains poorly understood. Here, we identify kinetically coupled domains that include post-translational modification and oncogenic mutation sites. We uncover networks of allosteric signaling pathways that provide fundamental insight into how the nucleosome communicates.

## 1. Introduction

The shape of biomolecules such as proteins and nucleic acids is intimately linked to their function. Proteins often undergo conformational changes upon binding nucleic acids to perform tasks including catalysis^1, 2^, signaling^3^, and transport^4^. Similarly, nucleic acids dynamically reorganize to modulate DNA accessibility^5^ or RNA structure during translation^6^. These conformational shifts are frequently triggered by specific ligands or environmental cues, allowing molecules to function as biochemical sensors^7^. For instance, enzymes may shift between active and inactive states in response to cellular energy levels. At the same time, transcription factors can bind to double-stranded DNA in the nucleosome, altering gene expression patterns. Traditionally, allostery has been associated with structural transitions in response to external stimuli; however, the concept of dynamic allostery has expanded this view by focusing on changes in flexibility and disorder within the same overall conformational space^8^. Such subtle dynamic modulations—such as fluctuations in dihedral angles or the transient breaking of hydrogen bonds—can propagate signals across a molecule without necessitating large-scale structural rearrangements, thus providing a mechanistic basis for long-range communication^9^.

A range of computational and theoretical methods have been developed to characterize correlated structural changes in biomolecules, spanning from elastic network models for proteins^10^ to base-pair-level descriptors of DNA deformability^11^. Additional approaches include principal component analysis (PCA), which captures dominant modes of motion in biomolecules^12^, dynamical network models^13^ that map residue communication pathways and capture local coupling dynamics, including for DNA base pair orientations^14^, and graph-theoretical frameworks that represent molecular structures as networks to identify key topological features underpinning allostery and conformational dynamics^15^. Time-lagged independent component analysis (tICA) identifies slow collective motions and potential allosteric pathways by capturing time-correlated features in molecular dynamics trajectories^16^. tICA can serve as a basis for the development of Markov State Models (MSMs) that capture long-time dynamics; for example, our laboratory has applied MSMs to map the long-timescale conformational dynamics of histone tails, revealing metastable states and transition pathways that may be subtly altered with post-translational modifications (PTMs)^17^. MSMs have also been used in combination with coarse-grained models to characterize the dynamic landscape of nucleosome assembly^18^. Additionally, methods in machine learning, such as diffusion maps, have been employed to characterize the translocational and rotational motion of complex polymers, including DNA within the nucleosome^19^.

To track changes in the molecular degrees of freedom, concepts like the mutual information theory have been used, which measures the amount of information transmission between two residues in a system, regardless of the functional form of their relationship^20^. While heterogeneity in structure can be addressed using parameters such as normalized mutual information^21^, heterogeneities in dynamics can be further understood by adapting methods from glassy systems^22, 23^. By combining mutual-information measures of structural co-variation with glassy-systems tools that capture transient, locally heterogeneous motions, we get a unified framework to map how allosteric signals are transmitted. Together, these approaches help bridge structural and dynamic heterogeneities by identifying both static correlations and temporally coordinated motions across biomolecular systems. Graph models, in which specific atoms in amino acids and nucleotides are depicted as nodes and their connections as edges, have also been used to trace the route between sites of allosteric induction and the site of the downstream effector^2, 24, 25^. The resulting node and edge connection generates a weighted graph system whose substructures of highly correlated residues can be determined with the Girvan-Newman algorithm^26^. The shortest path (Floyd-Warshall algorithm) linking allosteric sites aids the understanding of information transmission in the constructed network^25, 27^. Indeed, regions of proteins or DNA may behave in a more correlated or more liquid-like way when incorporating key features that are sometimes ignored in the approaches mentioned above.

Here, we focus on side-chain dihedral angles, a feature not commonly incorporated in previously discussed approaches. However, the timing transitions of changes in the side-chain dihedral angle can be used to characterize how distant regions in proteins communicate, as was shown with Correlation of All Rotameric and Dynamical States (CARDS)^21^. We next extend a similar approach to characterize the interface between protein-DNA, incorporating the base-sugar torsion angle of the DNA, which is known to be coupled to distinct transitions in DNA structure^28, 29^. Various biophysical techniques, including mass spectroscopy, nuclear magnetic resonance (NMR), and other computational techniques like molecular dynamics simulations, have been employed to study the allosteric properties of proteins and nucleic acids^30-34^. Flexible and interacting sites in proteins have been seen using highly sensitive hydrogen-deuterium exchange mass spectroscopy^35^. These techniques capture the dynamics of different domains of the molecule, examining dynamically correlated regions by measuring a specific degree of freedom.

The nucleosome core particle (NCP) is one of such allosteric complexes whose dynamics are affected by allosteric regulators, post-translational modifications, and interaction with transcription and remodelers. Molecular dynamics simulations revealed that histone variants macroH2A and an L1-loop mutant enhance nucleosome stability by strengthening dimer-dimer and histone-DNA interactions, reducing DNA breathing, and reinforcing compact conformations^36^. In contrast, H2A.Z introduces slight energetic changes that reconfigure allosteric communication networks, potentially modulating chromatin accessibility and regulatory response. Our laboratory has observed that acetylation of the H2B tail alters tail interaction with the double-stranded DNA in the NCP^37^. Also, a mutation in the histone core, including those implicated in certain cancers, has been shown to change the hydrogen bond dynamics in the core histones, indicating information is transmitted within the NCP^38^. Critical disruptions at nucleosome interaction interfaces have been seen to rewire chromatin regulation in histone-associated oncogenic systems (H2BE76K, H3K27M, H3K36M, and H3G34R), contributing to oncogenesis^39^. Nucleosome interaction histone-histone, histone–DNA, and histone–protein interfaces were rewired, affecting dynamics and changing nucleosome allostery. Several additional PTMs and mutations highlighted in our analysis occur at functionally critical residues identified through conditional activity mapping. For instance, H3R42—a methylation site within the DNA entry/exit latch region—exhibits high inter-residue conditional activity with other PTM sites such as H3K27 and H3K36, suggesting coordinated dynamics across the nucleosome. H2BE76, a recurrent oncogenic mutation site, shows strong coupling with H3R131 and H2AH31, indicating long-range communication between histone cores. These residues, found at the histone-DNA interface, in the acidic patch, or at the tetramer-dimer junction, form kinetically linked networks that are sensitive to both PTMs and mutation-induced perturbations, with potential implications for nucleosome stability and chromatin regulation.

Given the extensive network of allosterically linked PTM and mutation sites, it is critical to adopt a framework that captures the underlying kinetic relationships driving these long-range interactions. Entropic properties, such as distances and dihedral angles, derived from molecular dynamics simulation trajectories, were utilized by Vögele and coworkers in their software package, Python ENSemble Analysis (PENSA), to comprehensively investigate changes in biomolecular conformational ensembles^40^. Lin et al.^9^ proposed the use of kinetic properties in their time-related kinetic correlation, which they termed conditional activity, to trace the correlation between domains in Catabolite activator protein (CAP)^9^. Lin (2016) uniquely reveals a hidden allosteric link in CAP, a directional, dynamic coupling from the cAMP-binding site of one monomer to the DNA-binding site of the other, undetectable by mutual information or RMSD-based structural analysis. In a sister study, Manley and Lin revealed that Ras isoforms use distinct combinations of kinetic (timing-based) and thermodynamic (entropy-based) allosteric pathways centered on Switch I and II to regulate effector interactions and catalytic activity, explaining isoform-specific functional differences^41^. To identify residues whose changes in degree of freedom are significant in the overall dynamics of the NCP and reveal kinetically correlated domains in the system, we introduce the open-source Python library *CONDACT*, based on the concept of conditional activity proposed by Lin et al^9^. *CONDACT*, coined from CONditional ACTivity, traces time-related changes of thermodynamic properties in the trajectory of long-scaled molecular dynamic simulation trajectories to deduce kinetically significant residues and domains and trace the route of information transmission between the site of allosteric induction and the site of effector response. The library is also sensitive to identifying long-range communication over nm distances in space.

## 2. Methods

### Conditional Activity

Molecular dynamics is characterized by changes in various degrees of freedom, which include dihedral angle, distances, electrostatic contacts, among others. While using molecular dynamics simulation to sample the potential energy surface of biomolecular systems, different residues and domains adopt different states at any given time during the system’s dynamics. The dynamic relationship among different parts of heterogeneous systems, like proteins, has been computed using different correlation tools. Mutual information (MI) theory has been widely used to understand the correlation between two amino acids X and Y in protein systems. The MI of an observable degree of freedom of X and Y is defined as the sum of the entropies of X and Y minus the combined entropy of the complex X and Y^42, 43^, as follows *MI*(*X, Y*) = ∑_*x*∈*X*_ *P*(*X*) *ln*[*P*(*X*)] + ∑_*y*∈*Y*_ *P*(*Y*) *ln*[*P*(*Y*)] − ∑_*x*∈*X*_ ∑_*y*∈*Y*_ *P*(*X, Y*) *ln*[*P*(*X, Y*)]. Here, *P*(*X*), *P*(*Y*), and *P*(*X, Y*) are the probabilities of *X, Y* and *X,Y* respectively. The MI, therefore, gives the amount of thermodynamic or entropic information that X shares with Y.

However, some residues or domains in some biological systems exhibit large-scale kinetic correlation with or without noticeable physical changes in their entropic or thermodynamic properties. The correlation of these residues can also be evaluated using kinetic information, termed conditional activity^9, 41^. Degrees of freedom like distance, angle, electrostatic contacts, and backbone or side-chain dihedral angle change states, and the changes in these states serve as a kinetic clock to track kinetically associated residues/domains. These degrees of freedom are usually characterized as being in one finite state at any point in time during a molecular dynamics simulation. The time at which these degrees of freedom of a residue, X, change from one finite state to another is termed the transition time of the amino acid (*T*_*x*_). If two residues *X* and *Y* have a degree of freedom that changes among different finite states, the *i*^th^ transition time of *X* or *Y* during a simulation denoted as *T*(*X, i*) and *T*(*Y, i*), respectively, is the time at which the degree of freedom of *X* or *Y* change from one finite state to another finite state. The waiting time of the residue *X*, denoted as *W*(*X, t*), is the time interval between the time of the *i*^th^ transition and the (*i+*1)^th^ transition of X as follows, *W*(*X, t*) = *T*(*X, i* + 1) − *T*(*X, i*). The transition and waiting times are independent of the state of the degree of freedom of interest but depend on the time of transition of that degree of freedom. The conditional activity is related to the persistence time of specific residues and the exchange time between two residues.

The mean persistence time is the average time a degree of freedom of a residue stays in a specific finite state during a simulation. It is therefore the average waiting time starting from a random time *t* until the next transition of the degree of freedom of X. To accurately compute the persistence time of an amino acid, the number of transitions by the degree of freedom must be far greater than 1 (N >>1). Therefore, the conditional activity between residues is best determined for a long simulation trajectory with an observation time *τ* ≡ T(X, N(X)) in which there are multiple transitions of the degree of freedom. The mean persistence time of *X, τ*_*p*_[*X*], is directly proportional to the mean squared waiting times of *X* calculated from the corresponding list of transition times as follows, 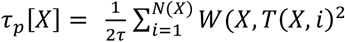. The mean squared waiting time is the product of that waiting time and the probability of selecting that waiting time. The probability of selecting a specific waiting time is proportional to that same waiting time. The mean exchange time of the degree of freedom of the residue *X* following a transition of another amino acid Y (*τ*_*p*_[*X*][*Y*]) is the waiting time for *X* to undergo a transition after an (*i+1*)^th^ transition in *Y*. The exchange time of *X* after a transition in *Y* is proportional to the probability of the waiting time between the *i*^th^ and (*i+1*)^th^ transitions of *Y* defined as 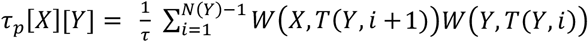. The conditional activity of X with respect to Y (A[X][Y]) is the negative logarithm of the ratio of the exchange time of X after a transition in Y (*τ*_*p*_[*X*][*Y*]) to the persistence time of X (*τ*_p_ [*X*]) as shown below, 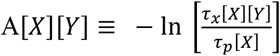.

If the degree of freedom of a residue *X* is independent of another residue Y, then the conditional activity of *X* on *Y* is equal to zero (A[*X*][*Y*] = 0). The more positive the conditional activity (A[*X*][*Y*] > 0), the higher the probability that a transition in *X* leads to a corresponding transition in *Y*. However, if A[*X*][*Y*] < 0, a transition in *X* is less likely to promote a corresponding transition in *Y*. Furthermore, the conditional activities of the degree of freedom of two amino acids are not commutative, i.e, A[*X*][*Y*] ≠ A[*Y*][*X*]. The diagonal of the conditional activity matrix is the conditional activity of a residue on itself (A[*X*][*X*]), and this represents the dynamical memory of that residue. If A[*X*][*X*] ≈ 0, then the successive waiting times of X are independent of the history of *X*, hence, the transitions of X are random following a Poisson distribution. The dynamical memory of a specific residue X is not interaction-specific and therefore doesn’t deduce the direction of interaction between X and another residue Y (X to Y). The dynamical memory only identifies residues whose side-chain dynamics are statistically significant. The direction of interaction between a residue X and another residue Y (A[*X*][*Y*]) can be obtained from the off-diagonal entries of the condition activity matrix, in which the more positive *A[X][Y]*, the higher the probability that the sidechain dynamics of X influences the sidechain dynamics of Y. The kinetically connected domain can also be traced from the conditional activity matrix. The principal eigenvector of the symmetrized conditional activity matrix shows the different domains of the system that are dynamically connected. This shows domains with the largest mode of correlated fluctuations, identifying residues with the highest contributions to these fluctuations.

### State Assignment to Degree of Freedom

We apply our conditional activity module to four systems: two enzymes and two nucleosome core particle (NCP) systems. Lysozyme (PDB ID: 1AKI)^44^ and the third domain of synaptic protein PSD-95 (PDZ3 with PDB ID: 1BFE)^45^ are the enzyme systems, while the nucleosome core particles (NCPs) contain the alpha satellite sequence (PDB ID: 1KX5)^46^ and the Widom-601 sequence (PDB ID: 3LZ0)^47^. The foundational 2.8 Å X-ray structure of the nucleosome core particle established the histone octamer architecture and DNA path, providing a structural baseline for mechanistic studies of dynamics^48^. Details of the system parameterization and simulation parameters can be found in the supporting material. The enzyme systems were each run for up to 3 µs, while each NCP system was run for 6 µs. The lysozyme and PSD-95 systems are analyzed to validate our CONDACT code and compare with the previous results reported in Lin (2016)^9^.

To understand the conditional activity of both proteins and DNA in the nucleosome core particle (NCP), dihedral angles are utilized as a measure of the degree of freedom. The large size of the NCP encourages slow dynamics of the systems, preserving the secondary structure of the core histones and double-stranded DNA. To this end, side chain dihedral angles (***χ***1) are used as the degree of freedom for the proteins (**Figure 1a**). In contrast, the base-sugar nucleoside dihedral angles (O4^I^-C1-N9-C4 for purine and O4^I^-C1-N1-C2 for pyrimidine) are used as the degree of freedom for the double-stranded DNA (**Figure 1 b-c**). The side chains of amino acids and the base-sugar backbone of nucleoside can adopt different states driven by short-range or long-range interactions of the residues. In essence, amino acids glycine and alanine, devoid of a side chain dihedral angle (***χ***1), are ignored in this study. The secondary angles of all negative dihedral angles are utilized to make a 0-360° range. The probability density of the dihedral angles of all amino acids and DNA was made (**Figure 1d-e**), and states are assigned to each peak. The protein dihedral angles are assigned three states (X, Y, and Z) while the DNA base-sugar nucleoside dihedral angles are assigned two states (S and A for syn and anti-conformations, respectively) as shown in **Figure 1b-c**. For amino acids, 0°≤ **θ**< 120° was assigned state X, 120°≤ **θ**< 240° was state Y, while 240°≤ **θ**< 360° was state Z. Similarly, for the nucleotides in the double-stranded DNA, 0°≤ **θ**< 130° and 330°≤ **θ**< 360° were state S and 130°≤ **θ**< 330° was state A. To ensure robust analysis, only residues with ten or more transitions are included in the conditional activity analysis because different states of the degree of freedom have been efficiently sampled.

**Figure 1:**
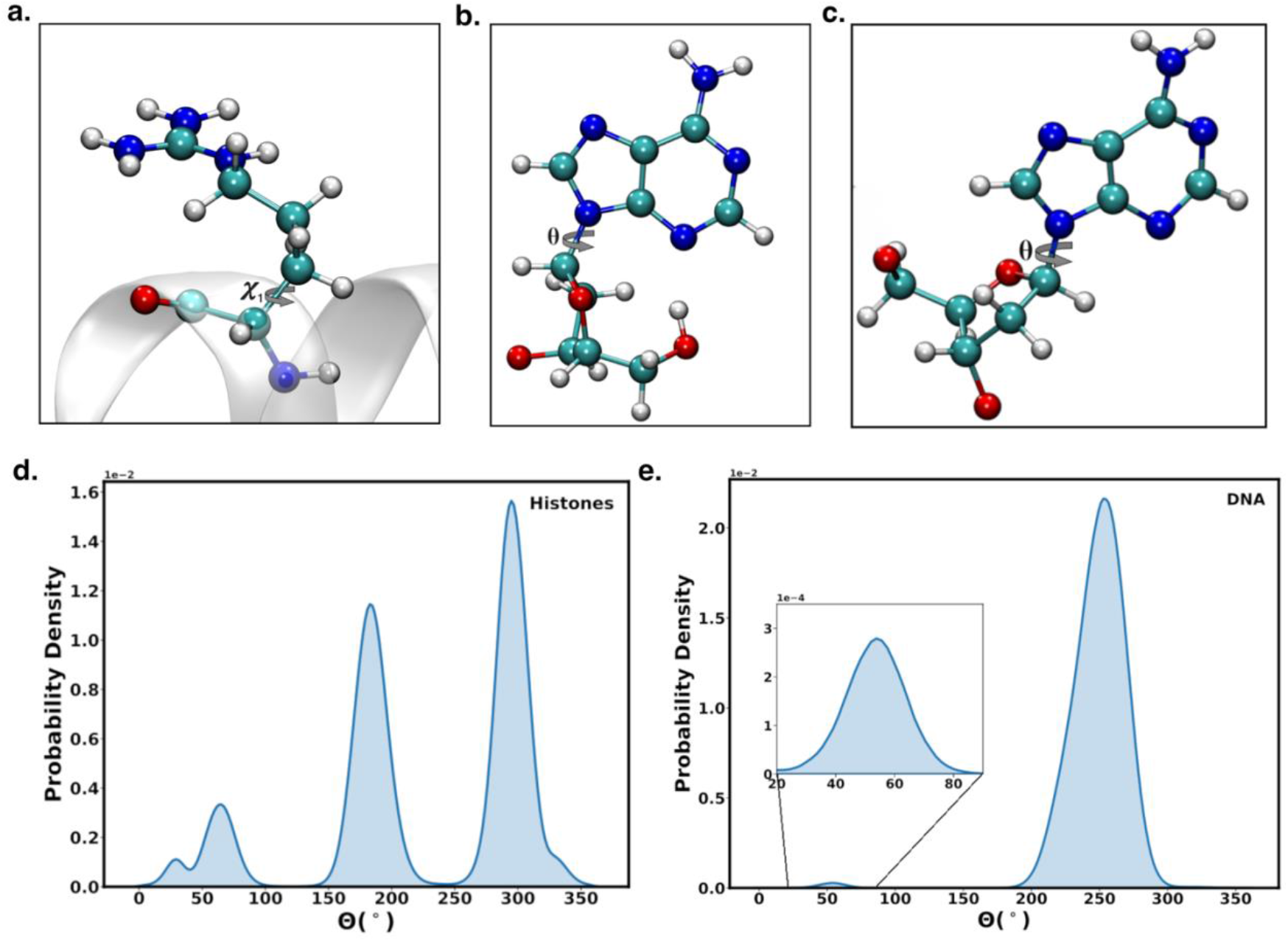
Dynamics of the dihedral angle of amino acid sidechain in protein and base-sugar dihedral angle in DNA is used as degree of freedom (a) First sidechain dihedral angle for proteins (***χ***^1^). The base-sugar dihedral angle (**θ**) for DNA in the (b) Syn conformation and the (c) Anti Conformation. (d) Probability density distribution of the sidechain dihedral angles for amino acids. The probability density shows three distinct peaks representing three states X, Y, and Z. (e) Probability density distribution of the base-sugar dihedral angles for DNA. The DNA probability density distribution of the dihedral angle reveals two defined peaks, which represent the syn (S) state and the anti (A) state.

## 3. Results

### Conditional Activity revealed residues with high dynamic memory

The conditional activity of all systems reveals amino acids or nucleotide residues in the enzyme and NCP systems, which are characterized by a high dynamical memory. The dynamics of residues with high dynamical memory do not follow a Poisson distribution; hence, their transitions are not random, or successive transitions depend on previous dynamics. For the enzyme systems, the residues with the highest dynamical memory are in the active site for lysozyme and in both the active and allosteric sites for PDZ3 (**Figure S1a-b, SI Text 1.3**), consistent with previously reported results by Lin et al^9^. For the NCPs, the dynamical memory for the degree of freedom is heterogeneously distributed throughout the systems (**Figures 2a-b**), revealing statistically significant dynamics of amino acid sidechains in the histone and base-sugar dihedral angle of the double-stranded DNA. These residues are in different domains on the histones, including the alpha-1 (α1) helix, alpha-2 (α2), and Loop1 (L1) of Histone H4, the H3-αN helix, the acidic patch, and the SHL-2 and SHL-4 regions in both the 1KX5 and/or the 3LZ0 systems. Single-molecule force spectroscopy revealed that nucleosomes unwrap asymmetrically from entry/exit sites and progress through SHL regions under tension, contextualizing the elevated dynamical memory we observe at SHL-2 and SHL-4^49^. It is worth noting that all histone subunits in the NCP had residues with high dynamical memory located in the core, tail, histone-histone interface, and histone-DNA interface (**Table 1**).

**Table 1:** Percentage of Residues with Dynamical Memory more than 4 in histone/NCP Domains.

|  | NCP Tails | NCP Core | H3 | H4 | H2A | H2B |
| --- | --- | --- | --- | --- | --- | --- |
| **1KX5** | 3.50 | 16.90 | 12.67 | 16.03 | 18.81 | 11.27 |
| **3LZ0** | 1.52 | 18.69 | 12.27 | 15.72 | 19.57 | 9.80 |

**Note:** Values reported in %

**Figure 2:**
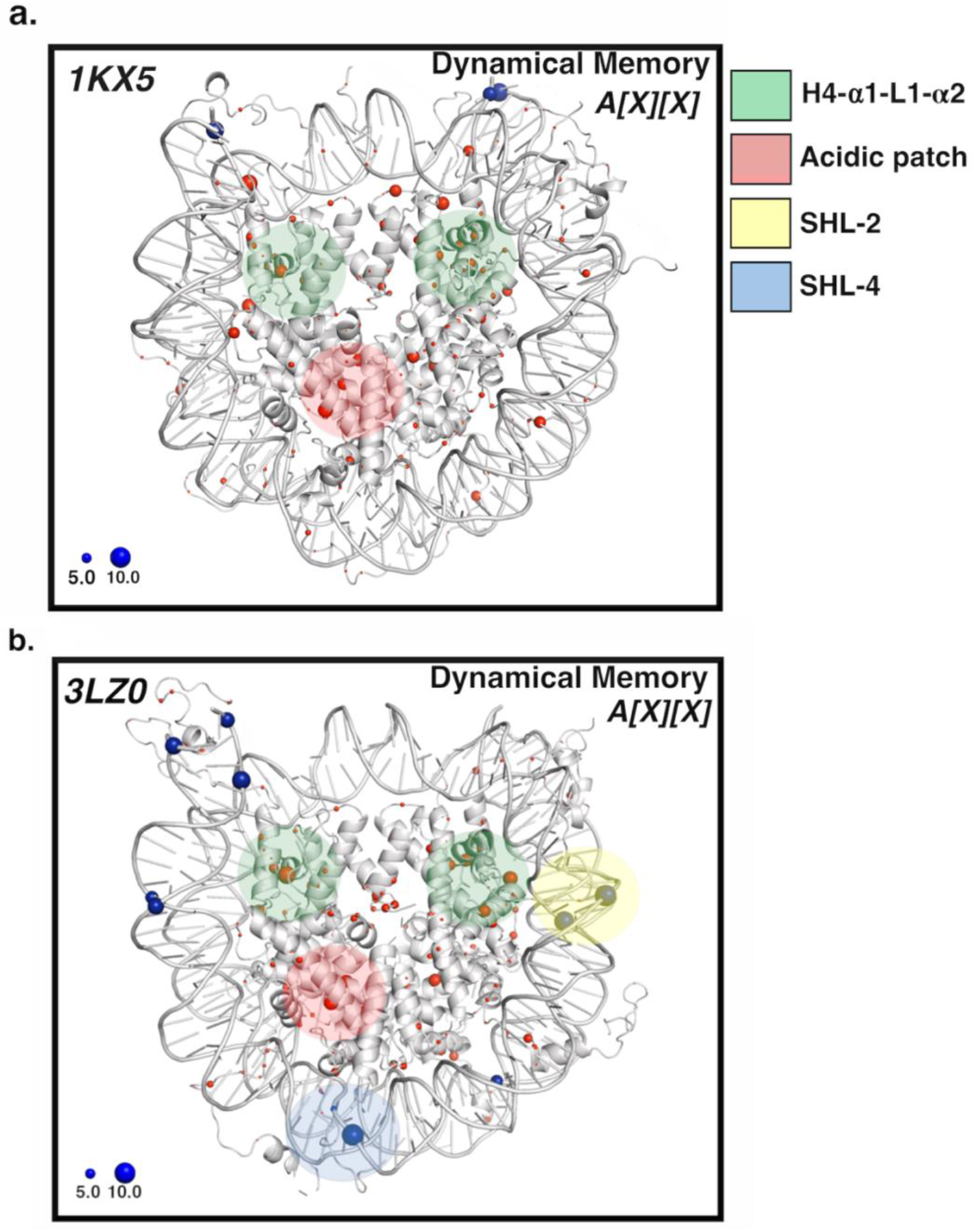
Dynamical memory (A[X][X]) of charged residues in the NCP for (a) alpha satellite sequence (1KX5) and (b) Widom-601 sequence (3LZ0). The sphere sizes are proportional to the dynamical memory. Residues with the higher dynamical memory at the H4-α1, H4-α2, and H4-L1 domains are in the green shaded region, the acidic patch residues are in the red shaded region, the DNA SHL-2 region is in yellow, while the SHL-4 region is in blue. The residues in these regions modulate the binding of nucleosome binding chaperones or remodelers, and their mutations have been seen in some cancer types, like breast cancer and colorectal cancer.

In the 1KX5 system, residues on both copies of the H4-α1, H4-α2, H4-L1, and the acidic patch show a very high dynamical memory (**Figure 2a**). The residues H4D24, H4R35, H4R36, and H4R40 on H4-α1, and H4K44 on H4-L1 show a high dynamical memory. Also, H4E53 and H4R55 on the α2 helix of H4.1 and H3R42 on the αN helix show statistically significant dynamics of their side-chain dihedrals. In the same light, H2AE56, H2AE64, and H2BD68, located in the acidic patch, have high dynamical memory.

The 3LZO system also has some identical dynamically active residues as the 1KX5 system. Both copies of histone H4 showed dynamically active residues on their α1 (H4R36 and H4R40) and α2 (H4E53 and H4R55) helices. Also, H3R63 on the α1-helix of both copies of histone H3 was dynamically active. The conditional activity of acidic patch residues in the 3LZ0 systems reveals that the transitions of H2AE56, H2AE64, H2AD91, H2BD68, and H2BE113 are not memoryless, as the dynamical memory was high. However, unlike the 1KX5 system, some nucleotides in the double-stranded DNA in the 3LZ0 systems, which are distant from the DNA terminals, show statistically significantly high dynamical memory. For example, Guanine 54 (5^I^-3^I^ strand) and guanine 93 (3^I^-5^I^ strand) in the SHL-2 region and guanine 32 (5^I^-3^I^ strand) in the SHL-4 region showed high dynamical memory, which might indicate the residues are slightly stressed when flipping between the Syn and Anti states.

The residues on the H4-α1 helix have been implicated to interact with nucleosome remodelers and histone binding proteins. H4D24 has been seen to modulate the binding of chromatin remodelers like Chd1, disrupting the methylation of H4R20 by Suv4-20 homolog 1 (Set8/Suv4-20h1)^50^. Cryo-EM structures of Chd1 bound to nucleosomes show how remodelers engage the H4 tail/acidic-patch surfaces to stabilize sliding intermediates, aligning with the kinetically active H4/acidic-patch residues we detect^51^. The histone chaperone retinoblastoma-associated protein RbAp46 interacts with H4-α1 helix with H4R35, H4R36, and H4R40 actively involved in its binding^52^, while H4K44 has been seen to facilitate chromatin accessibility^53^. Residues in the acidic patch of the nucleosome not only regulate binding of the nucleosome to some chaperones like FACT (in yeast), but their mutations have been implicated in different cancer types like colorectal cancer, head and neck cancer, and breast cancer^54, 55^. The SHL-2 and SHL-4 regions are the sites where histone H4 N-terminal tail interacts with the DNA, or the sites at which pioneer transcription factors like Sox2 bind with the double-stranded DNA^56-58^.

To deduce the direction of association between two residues X and Y, the off-diagonal entries of the conditional activity matrix are used. The nucleosome is a highly allosteric complex with large-scale communication among residues in the histone and DNA. These pairs of communication is not only found in the histone core but also at the histone-DNA interface (**Figures 3a-b** and **S2c-d**). The amino acids with the highest inter-residue conditional activities include sites of oncogenic mutations and sites of post-translational modifications (**Table 2, SI Table 1**). Residues like H3R42, H3R131, H4R92, and H2BE76 were seen to have high inter-residue conditional activities affecting the side chain dynamics of multiple residues, both on the identical and other histone subunits.

**Table 2:** Inter-residue Conditional Activity greater than 6 of Residues at Oncogenic and Post-Translational Modification Sites.

| Residue-X | Location | Mut-X | PTM-X | Residue-Y | Location | Mut-Y | PTM-Y | A[X][Y] |
| --- | --- | --- | --- | --- | --- | --- | --- | --- |
| H2AH31 | H2A-α1 helix | Yes | No | H2BE76 | H2B-α2 helix | Yes | No | 6.31 |
| H2AH31 | H2A-α1 helix | Yes | No | H3K27 | H3 N-terminal | Yes | Yes | 6.25 |
| H2BF70 | H2B α2 helix | Yes | No | H2BE76 | H2B-α2 helix | Yes | No | 6.08 |
| H2BF70 | H2B α2 helix | Yes | No | H3E105 | H3 α2 helix | Yes | No | 6.03 |
| H2BF70 | H2B α2 helix | Yes | No | H3K27 | H3 N-terminal | Yes | Yes | 6.29 |
| H3R131 | H3 C-terminal | Yes | No | H2BE76 | H2B-α2 helix | Yes | No | 6.03 |
| H3R42 | NCP latch | No | Yes | H2BE76 | H2B-α2 helix | Yes | No | 6.92 |
| H3R42 | NCP latch | No | Yes | H3R26 | H3 N-terminal | Yes | Yes | 7.51 |
| H3R42 | NCP latch | No | Yes | H3E73 | H3 α1 helix | Yes | No | 7.42 |
| H3R42 | NCP latch | No | Yes | H3K27 | H3 N-terminal | Yes | Yes | 6.80 |
| H3R42 | NCP latch | No | Yes | H3K36 | H3 N-terminal | Yes | Yes | 7.67 |

**Note:** Mut-X is the site of oncogenic mutations of X
PTM-X is the site of post-translational modification of X
Mut-Y is the site of oncogenic mutations of Y
PTM-Y is the site of post-translational modification of Y
A[X][Y] is the conditional activity of X on Y

**Figure 3:**
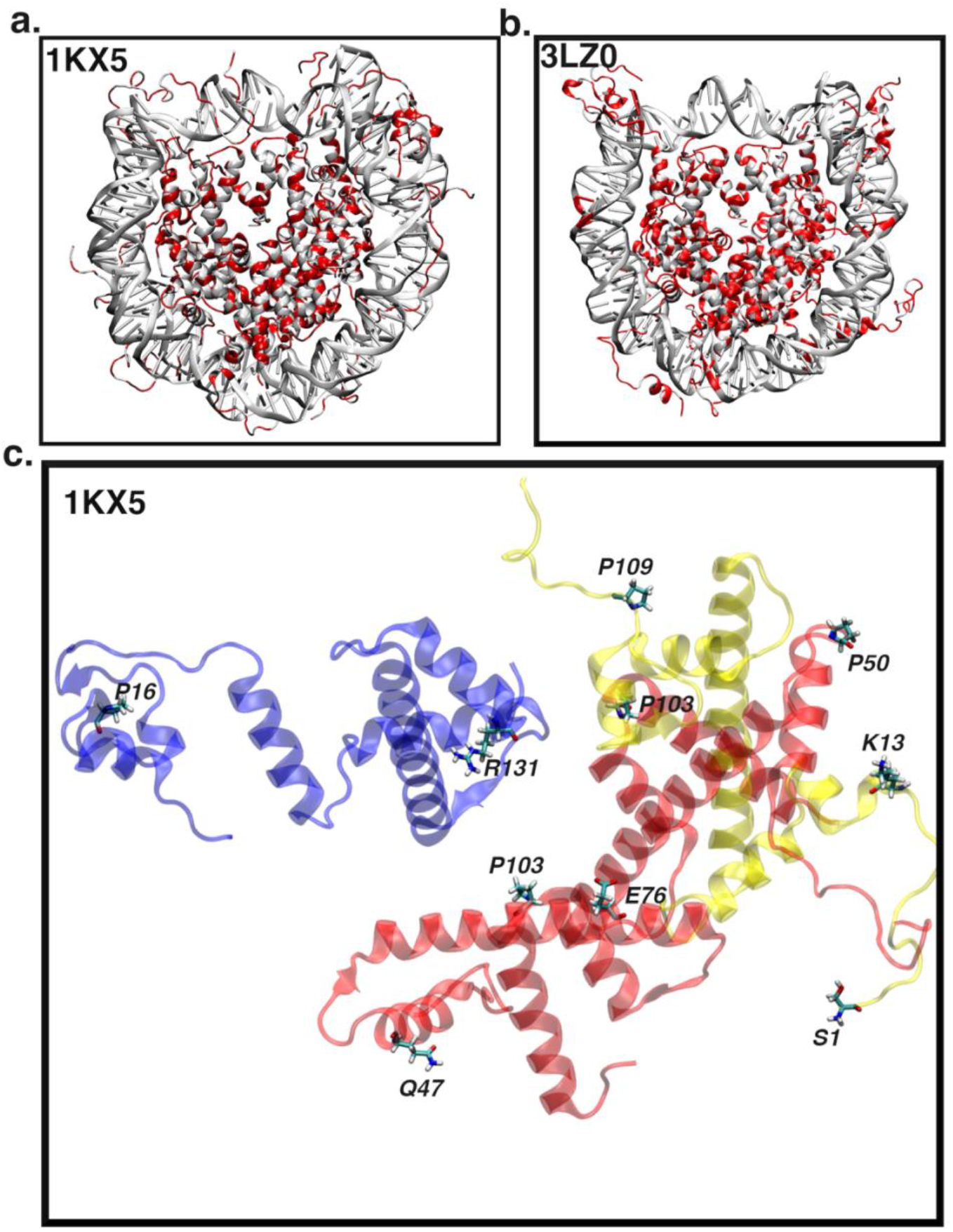
(a) Inter-residue conditional activity (*A[X][Y]*) between residues in the nucleosome. The core histone residues show highly correlated dynamics among themselves. (b) Positive inter-residue conditional activity between H3-R131 and other amino acids in the identical or other histone subunits. Histone H3 in blue, H2A in yellow, and H2B in red. Side-chain dynamics show long-range communication between H3-R131 and other amino acids, including H2B-E76 and H2A-K13.

The methylation of H3R42, a post-translational modification site at the DNA entry/exit region of the NCP, was seen to open the NCP by reducing the interaction between the double-stranded DNA and the histone core^59^. Armeev and colleagues described H3R42 and adjacent amino acids (H3H39 and H3R49) as an H3 latch, revealing they were the first barrier to DNA unwrapping in the NCP, hence very important for gene regulation^60^. Mechanistic models and cryo-EM suggest that remodelers drive nucleosomal motion via propagated DNA twist-defects, which intersect the latch/entry regions highlighted here^61^. The residues H4R92 and H2BE76 are core oncogenic sites, and their mutation have been previously seen by Onyema *et al*. to destabilize the histone core through changes in hydrogen bond dynamics^38^. **Figure 3c** shows the conditional activity of H3R131, an oncogenic site with other amino acids in the histone core. The sidechain dynamics of H3R131 via long-range interactions exhibit both intra- and inter-histone communications with other amino acid sidechains in the NCP. H3R131 has been suggested to contribute to the stability of the tetramer-dimer structure of the nucleosome core particle^62^. According to Ngubo and coworkers, H3R131 formed a hydrogen bond with H3Y99, H3D106, and H2AR99. Both intra- and inter-histone interactions by H3R131 can promote nucleosome stability.

### Principal Eigenvector Shows Communication between H2A-H2B Dimers

The nucleosome possesses dynamically connected domains, as indicated by its principal eigenvectors. The dynamically connected domains were nucleotide sequence-dependent, as shown in **Figure 4a-b** for the 1KX5 and 3LZ0 systems. The most dynamically connected domains are represented with red spheres, while the least are in green spheres. The domains with intermediate principal eigenvectors are in blue.

**Figure 4:**
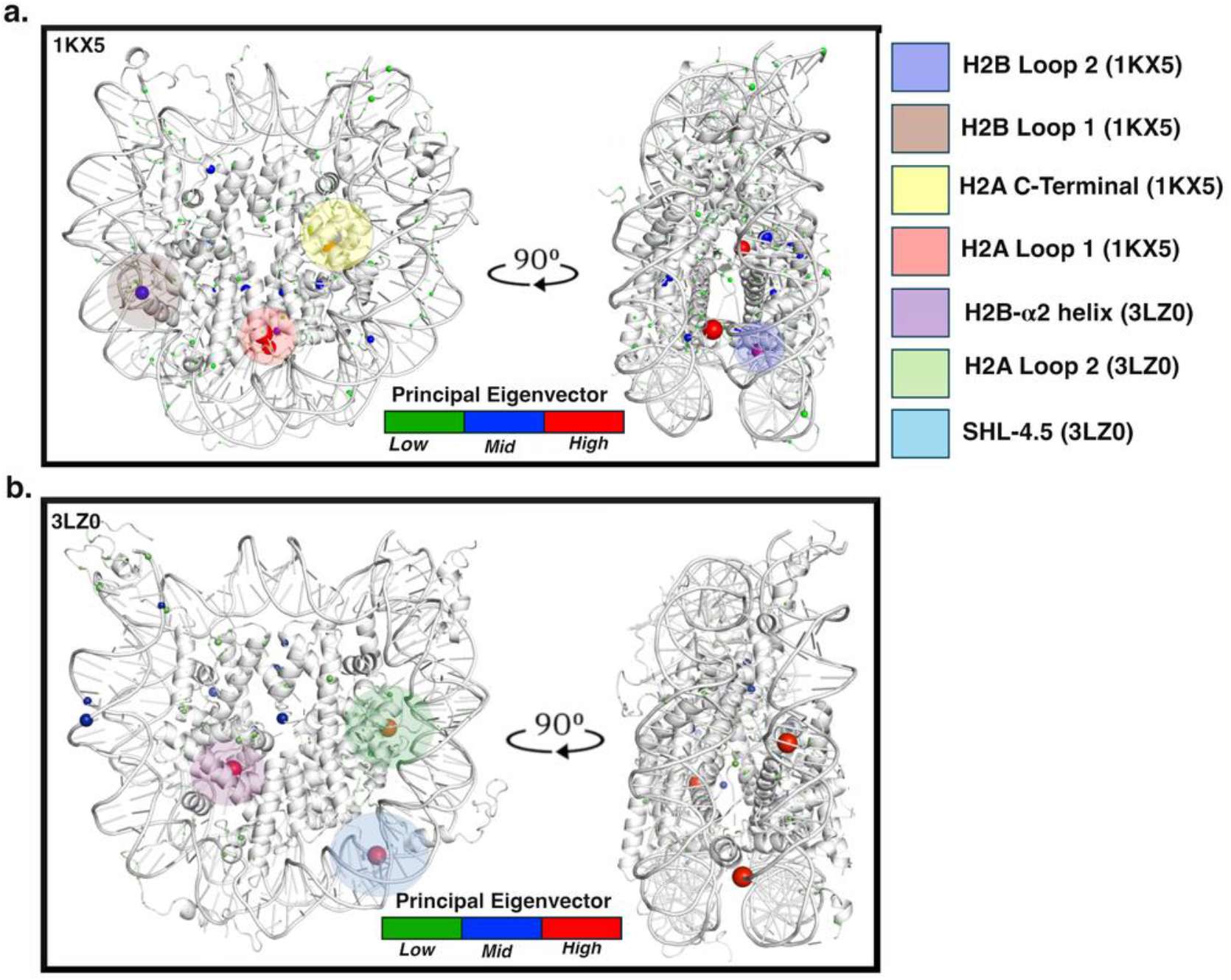
Dynamically correlated regions of the (a) 1KX5 systems and the (b) 3LZ0 system shown at 0° and 90° rotation. The sphere size is proportional to the contribution of the residue to the principal eigenvector calculated from the symmetrized conditional activity matrix. In the 1KX5 system, regions whose degrees of freedom are kinetically connected include the C-terminal tail (yellow-shaded region) of H2A, loop 1 of H2A (red-shaded region), loop 1 (navy blue-shaded region) of H2B, and loop 2 (brown-shaded region) of H2B. In the 3LZ0 system, the α2 helix of H2B (violet-shaded region), loop 2 of H2A (green-shaded region), and the SHL-4.5 region (light-blue shaded region) of the Widom-601 DNA are kinetically correlated.

In the 1KX5 system, there are connections in the dihedral angle between the histone H2A and H2B in the handshake region (**Figure S3a-b**) and between the two H2A-H2B dimers (**Figure 4a, Figure S3c**). H2AY39 and H2BH49 primarily promote the kinetically connected domains in the first copy of the H2A-H2B dimer in loop 1 of each histone. In contrast, in the second copy of the H2A-H2B dimer, H2AI102 on the C-terminal tail, H2AQ84 on the α3 helix, and H2BT88 on loop 2 promote the kinetic connections of those domains. Similarly, H2AI102 and H2AY39 are suggested to foster dimer-dimer communication. Furthermore, both H2A-H2B dimers are kinetically connected to H4T71 in the α2 helix and H4I46 in loop 1 (**Figure S3d**). In essence, there is a kinetic connection between the H3-H4 tetramer and both H2A-H2B dimers **(Table S2)**.

Besides the kinetically connected domains above in both NCP systems, the 3LZ0 also shows a kinetic connection between loop 2 of H2A, promoted by H2AI79, the α2-helix of H2B, promoted by H2BF65, and G-45 in the SHL-4.5 region (**Figure 4b, Figure S4)**. Histone-DNA interaction at the SHL-4.5 region strains the sugar-base dihedral angle in G-45 (3^I^-5^I^ strand) from 290.5° to 331.5°, disturbing the Watson-Crick hydrogen bond **(Figure S5)**. Remodeler complexes can actively distort nucleosomal DNA. For example, INO80 generates local DNA bulges during translocation, providing a structural route for the long-range kinetic couplings we observe across histone–DNA interfaces^63^. There is also a tetramer-dimer kinetic connection at the C-terminus of both H3 (H3R134) and H4 (H4F100), loop 2 and α1 helix of H4 (H4V81 and H4I26), and α3 helix on H3 (H3L126) in **Figure S4**.

### Conditional Activity Shows Long-range Correlated Dynamics

The conditional activity of residues in the nucleosome shows high sensitivity over long distances. The dynamic association between residues with respect to their separation in space is determined for the 1KX5 and 3LZ0 systems. The fraction of kinetically associated residues within a specific distance was the ratio of correlated residues with a conditional activity of at least 2 to all residues within that specific distance. The dynamical memory, which is the conditional activity of a residue against itself (with a separation of 0 nm), showed that about 25% of the residues in the nucleosome showed strong kinetic association. Inter-residue association between residues as the spatial distance increases showed that the conditional activity was sensitive to residues that were up to 7.5 nm apart in space in the nucleosome system (**Figure 5**). At least 6 % of residues that were 7.5 nm apart were seen to dynamically communicate. However, such communication could barely be seen for residues that were 8.0 nm apart in space.

**Figure 5:**
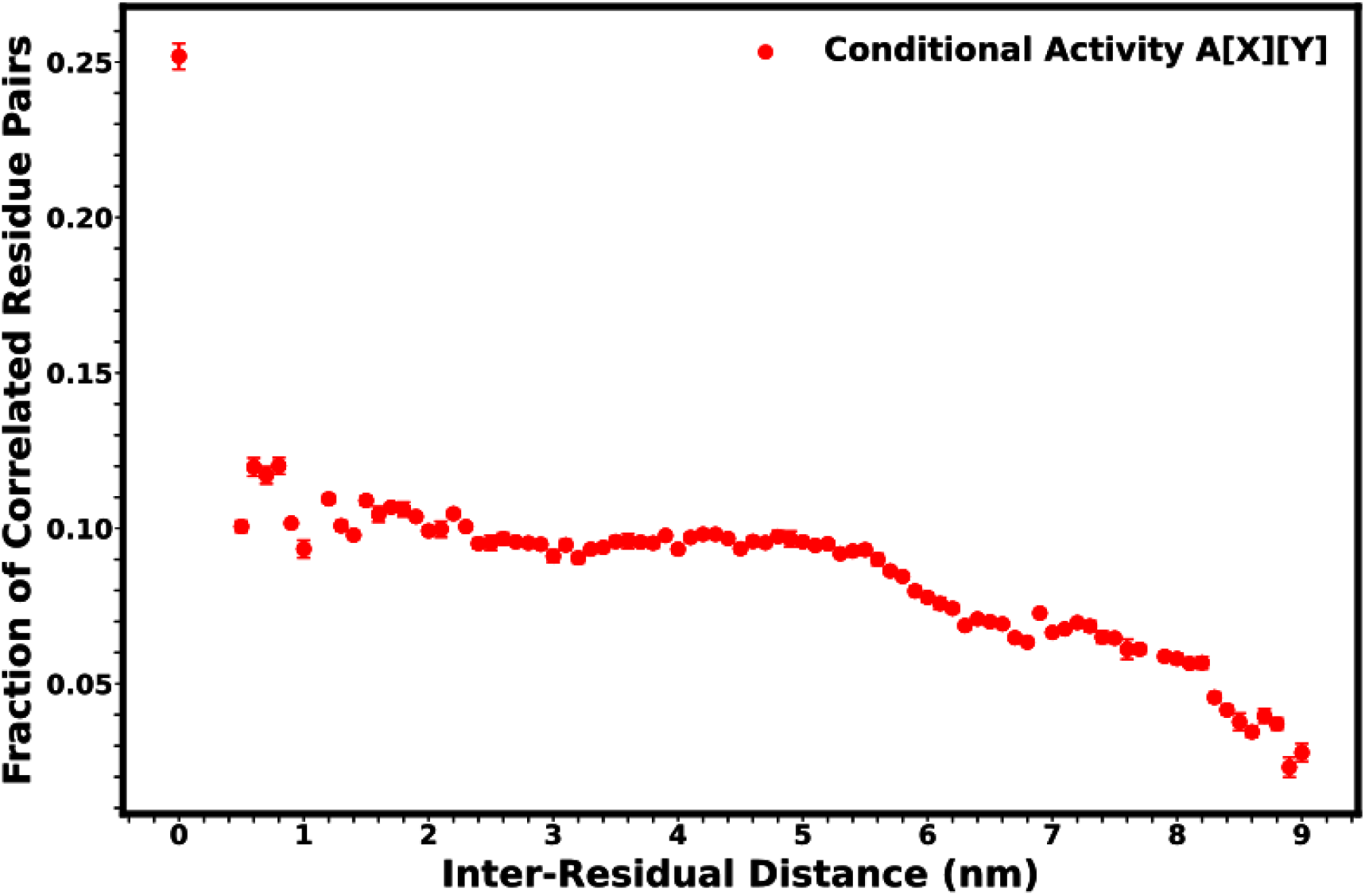
Comparison of the fraction of correlated dihedral angles against distance between two residues in the nucleosome systems. At least 6% of correlated residues show conditional activity of their dihedral angles at a 7.5 nm separation for the nucleosome system. The conditional activity drastically reduces after 8.0 nm. The error bar is the standard error of the mean obtained from replicate nucleosome systems.

## 4. Conclusion

Allosteric communication in biomolecular systems, particularly the nucleosome, plays a pivotal role in regulating cellular processes by coupling local perturbations with long-range conformational changes. Despite advancements in structural and thermodynamic analyses, understanding the kinetics of allosteric signaling, especially over extended spatial and temporal scales, remains a significant challenge. In this study, we introduce CONDACT (CONDitional ACTivity), a novel and open-source Python-based computational toolkit that quantifies kinetic correlations between molecular residues by evaluating time-resolved conditional activity. This approach leverages molecular dynamics (MD) simulations and builds upon Lin’s concept of conditional activity to map dynamically correlated domains and identify residues with high kinetic memory. Here, we apply CONDACT to four biological systems: two enzymes (lysozyme and PDZ3 domain of PSD-95) to validate the code, and two nucleosome core particles (alpha-satellite DNA and Widom 601 sequence NCPs). Using dihedral angle transitions as kinetic clocks, we assess the dynamical memory and inter-residue conditional activity, revealing residue-specific and domain-level communication patterns, including known oncogenic mutation sites and post-translational modification (PTM) hotspots. Our analysis highlights kinetically significant domains such as the H4-α1 helix, acidic patch, and DNA SHL-4.5 regions. The SHL-4.5 region is the site for DNA repair or remodeling, where DNA polymerase β can bind, carrying out one-nucleotide excision repair within intact chromatin^64^.

These findings not only confirm previously characterized interactions but also uncover new long-range kinetic correlations extending up to 7.5 nm, underscoring the sensitivity and range of CONDACT. The principal eigenvectors of the symmetrized conditional activity matrix further elucidate domain-level allosteric coupling, providing mechanistic insights into nucleosome plasticity and regulation. Adhireksan and colleagues saw that RAPTA-T (a ruthenium compound) binding to the acidic patch of the NCP induces conformational changes that allosterically enhance the formation of auranofin adducts at distant 36 Å H3 sites (H113), highlighting a novel paradigm for epigenetic targeting through coordinated modulation of chromatin structure^65^. Beyond local dynamics, heterochromatin proteins can reshape the nucleosome core and promote phase separation, offering an allosteric layer that meshes with our kinetic domain model^66^. Genome-wide sub-nucleosomal mapping further indicates distinct nucleosome folding modes, consistent with multiple kinetic states captured by our domain-level eigenvectors^67^. Overall, CONDACT bridges a critical gap in computational allostery by offering a kinetic-centric perspective and a scalable analytic platform that can be applied across diverse biomolecular systems. As a freely accessible tool, it lays the groundwork for future studies aimed at deciphering functional communication pathways, guiding drug discovery, and informing rational protein engineering design^68^.

Here, we identify crucial post-translational modification (PTM) sites and oncogenic mutations that show high inter-residue conditional activity, indicating strong dynamical coupling and functional significance. Key residues identified are within the acidic patch, such as H2AE56^69^, or on the H3 tail within the latch region identified by Armeev *et al*.^*60*^, such as H3R42. These sites are drug development hotspots, especially in the nucleosome, which is a challenging structure to target due to its tight packaging and dynamic regions like histone tails^70-72^. Traditional drug design often struggles with the nucleosome because many meaningful interactions are transient or disordered^71^. CONDACT overcomes this by focusing on how residues behave over time, rather than relying only on entropic structural correlations. Residues with high conditional activity may act as communication hubs within the nucleosome and could be key to stabilizing the structure in the presence of disease-related mutations. The binding of transcription factors or chromatin remodelers can change these activity patterns, disrupting or enhancing communication across the nucleosome. Understanding how these shifts occur may provide insight into how mutations alter nucleosome function and suggest ways to correct these changes with drugs or other synthetically designed biomolecules. This approach offers a new path for targeting nucleosome-related diseases such as cancer.

## Data Statement and Code Availability

Analysis codes are available on GitHub at https://github.com/CUNY-CSI-Loverde-Laboratory/conditional_activity.git. Trajectories are available on Zenodo.

## Acknowledgments

The NSF supported the work through the 2023-2024 MolSSI software fellowship, as outlined in the Fellowship Agreement Number 480718-19B75 and Prime Award No. CHE-2136142. NIH also supported this work through 1R15GM146228-01. Anton 2 computer time was provided by the Pittsburgh Supercomputing Center (PSC) through NIH grant R01GM116961. The Anton 2 machine at PSC was generously made available by D.E. Shaw Research. We thank the CUNY-HPCC, domiciled at the College of Staten Island, for their help in running our enzyme systems, and the CUNY Office of Research and Fellowships for awarding the CUNY Dissertation fellowship to A. C. O. We thank Prof. Milo Lin for discussions. A.C.O. is grateful for discussions with Dr. Phu Tang.

## Author Contributions

A.C.O. performed the simulations and developed the conditional activity module with active assistance from MolSSI software scientist C.D.; J.M. ensured code optimization and adherence to best practices in code development, and S.M.L. supervised the project.

## SI Description

Supporting material can be found online at x.

