## Supplementary Information for "Mapping Allosteric Communication in the Nucleosome with Conditional Activity"

**Supplementary Information Text**

### 1. System Modelling/Simulation

#### 1.1 System Modelling

Four systems were used for this study: Two enzyme systems and two NCPs. The enzyme systems were Lysozyme (1AKI.pbd) and PDZ3 (1BFE.pdb), while the NCPs were the alpha-satellite sequence (1KX5.pdb) and the Widom-601 sequence (3LZ0.pdb). AMBER force field was used to parameterize all the systems[1]. The amino acids of the protein in both enzymes and the nucleosome core particle (NCP) were parameterized with ff19SB[2], the DNA with OL15[3] , while OPC was used as the water model[4]. The $Mn^{2+}$ in the crystal structure of the NCP was substituted with $Mg^{2+}$. Monovalent ions ($Na^+$ and $Cl^-$) in all systems were parameterized using the method of Joung and Cheetham,[5] while the method of Li and Merz was used for divalent ions ($Mg^{2+}$ in the NCP)[6]. To properly model water-ion interactions, the Lennard-Jones parameters of monovalent and divalent ions were modified using the method of Kulkarni *et al*[7].

#### 1.2 Simulation

#### Lysozyme and PDZ3

The enzyme systems were minimized using the steepest descent algorithm for 10 ps on GROMACS. Each system was equilibrated for 120 ps under the NVT ensemble and 1 ns under NPT ensemble[8, 9]. Each simulation was run for 3 μs, saving the trajectory once every 10 ps. The Nose-Hoover thermostat[10] was used to maintain the temperature of the system at 310 K while the pressure was kept at 1 bar using the Parrinello-Rahman barostat[11]. The SHAKE algorithm was applied to freeze hydrogen bonds for the simulation to be run with a 2 fs timestep[12]. Both the Particle Mesh Ewald (PME)[13] algorithm and van der Waals cut-off were kept at 10.0 Å to compute long-range electrostatic interaction and non-polar interaction, respectively.

#### Nucleosome Core Particle

For the NCP systems, minimization with both conjugate gradients and steepest descent algorithms was done for 15 ps with Amber18[14]. The system was equilibrated in NVT ensemble for 20 ps to 310 K[8, 9]. Furthermore, NPT equilibration at 1 bar using the Berendsen barostat[15] was done for 100 ns, maintaining the temperature of the system at 310 K with a Langevin thermostat. A timestep of 2 fs was used for this equilibration as the SHAKE algorithm[12] was implemented to freeze hydrogen atoms. The van der Waals cut off was 12 Å while the Particle Mesh Ewald (PME)[13] algorithm was used to calculate electrostatic interactions. The equilibrated system was run for 6 μs on Anton-2[16] as described in our previous work.

#### Dynamical Memory in Enzyme Systems

The dynamical memory identified amino acid residues in the active site of lysozyme and in both active and allosteric sites of synaptic protein PSD-95 (PDZ3), whose dynamics are statistically significant to the activity of the enzymes (**Figure S1a-b**). These amino acids in the lysozyme systems included Glu35, Asp52, Lys33, Ser36, and Asn59, which have been implicated to be either important for binding or catalysis in lysozyme[17]. This was the same with the PDZ3 systems in which Phe20, Asp43, and Tyr92 at the active and allosteric site had the highest dynamical memory, hence involved in the enzyme activity of PDZ3. The location of these amino acids is similar to helices, β-sheets, and loops, which have been proposed to be involved in binding or allosteric regulation[18]. The off-diagonal values reveal the inter-residue conditional activity and depict which direction of association between any two residues (**Figure S2a-b)**. A positive Tyr92-Pro3 and Phe20-Pro3 conditional activities (A[Y92][P3] = 9.336; A[F20][P3] = 8.263) suggest that the binding and allosteric sites of PDZ3 might be communicating with the N-terminal tail of the enzyme.

The principal eigenvector of the symmetrized conditional activity matrix for the PDZ3 system showed that the αA and αC helices and the tail of the βB strand are dynamically connected. Also, the binding site domains of lysozyme are dynamically connected (**Figure S1c-d**).

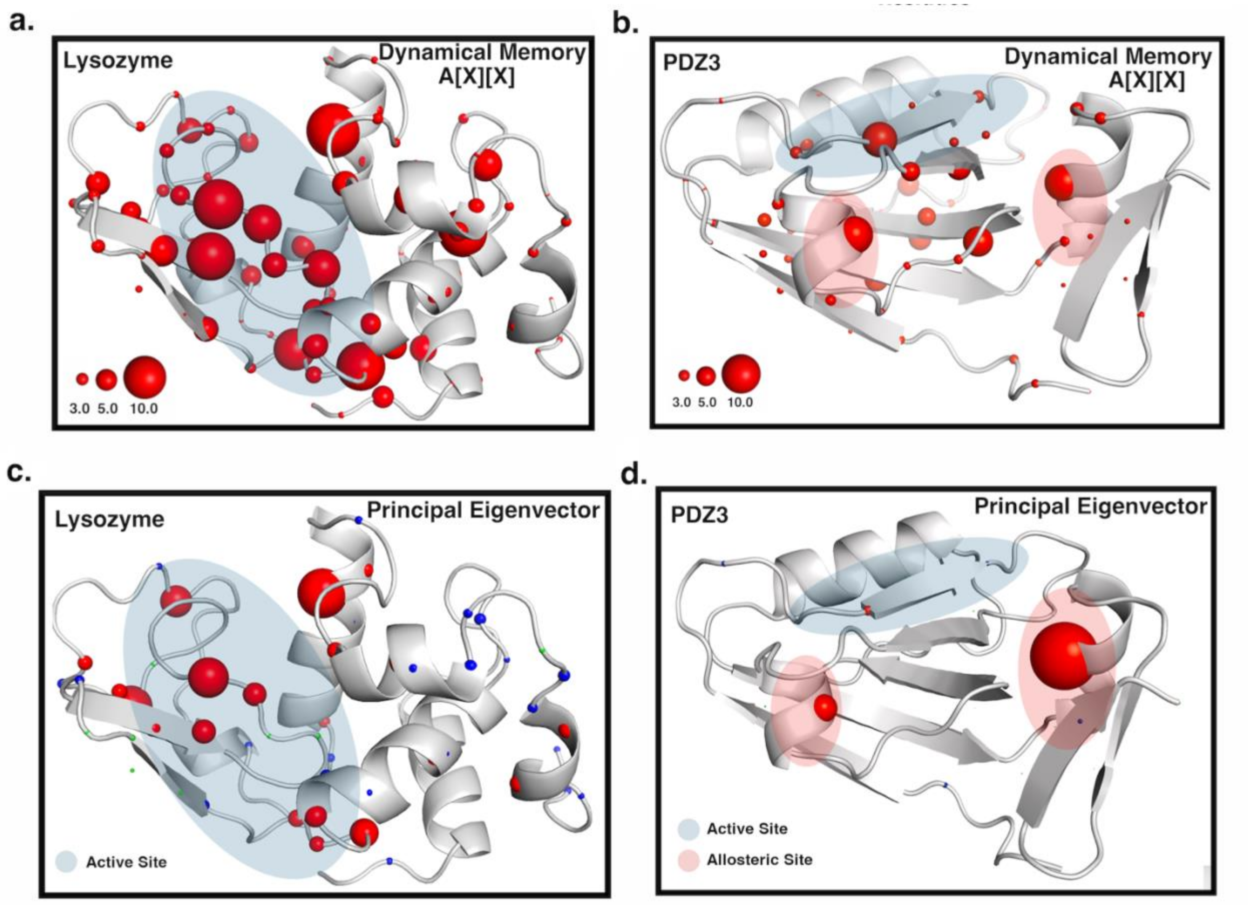


**Suppl. Figure 1**: Dynamical memory of all amino acids in (a) Lysozyme (b) PDZ3. The dynamical memory of the degree of freedom of amino acids at the active site (in blue) and allosteric site (in pink) is high; hence, their dynamics are statistically significant for enzyme activity. Principal eigenvector for (c) Lysozyme (d) PDZ3 reveals the regions with connected dynamics. The allosteric protein PDZ3 shows a sharp connection in dynamics among the allosteric αA and αC helices with the carboxylate binding loop, which interacts with ligands. Lysozyme shows a strong dynamic connection of helices and loops at the active site of the enzyme

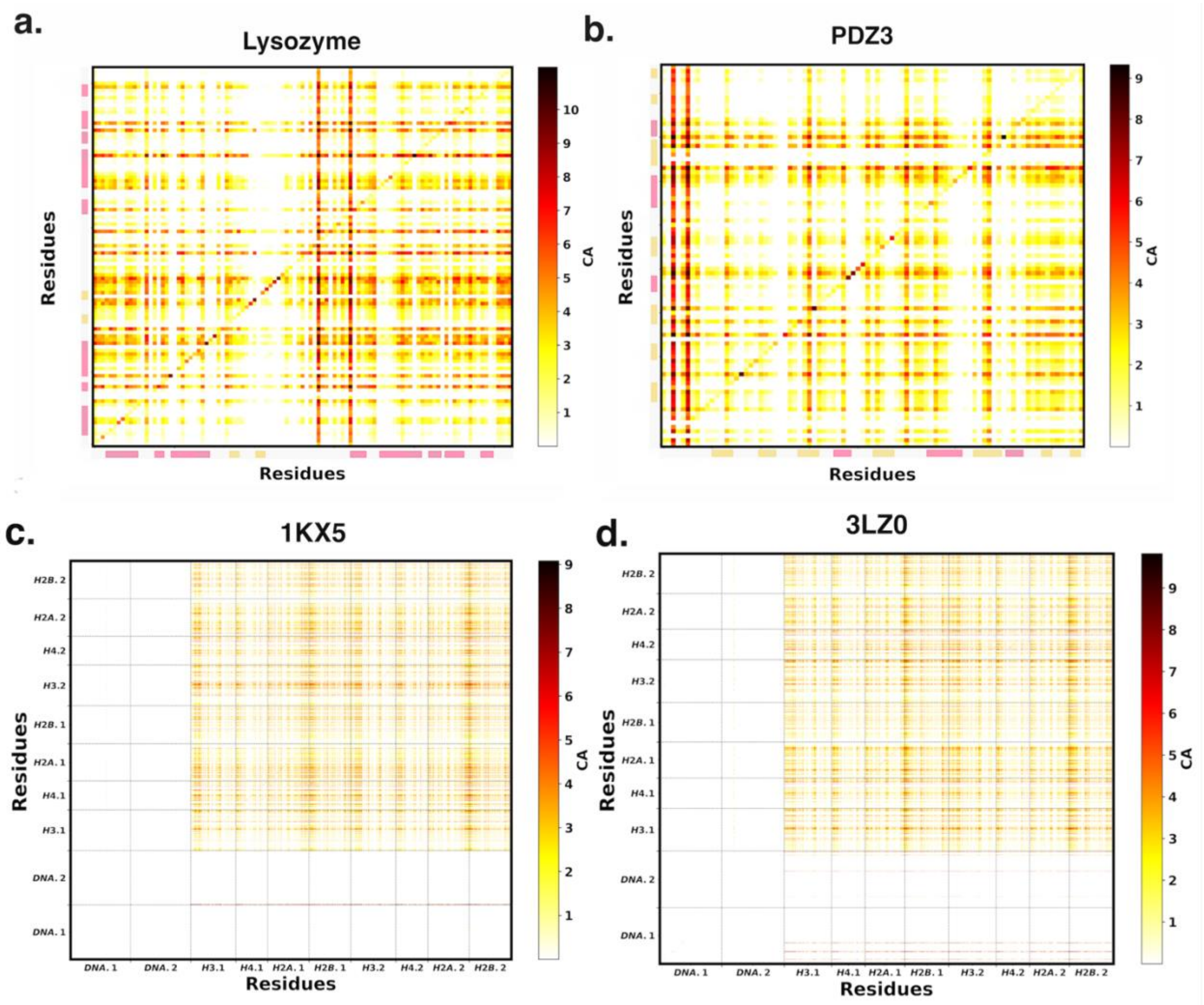


**Suppl. Figure 2**: Conditional activity of two enzymes; Lysozyme (1AKI.pdb) and Synaptic Protein PSD-95, PDZ3 (1BFE.pdb). Inter residue conditional activity heatmap for (a) Lysozyme (b) PDZ3. Pink rectangles depict α-helices, yellow rectangle **β**-sheets while interconnecting spaces are coils. Inter residue conditional activity heatmap for (a) 1KX5 (b) 3LZ0. High inter-residue conditional activity seen in the histone core while the double stranded DNA has little of no dynamics after 6 $\boldsymbol{\mu}$s. High conditional activity in the histone core underscores amino acid communication is important for nucleosome dynamics.

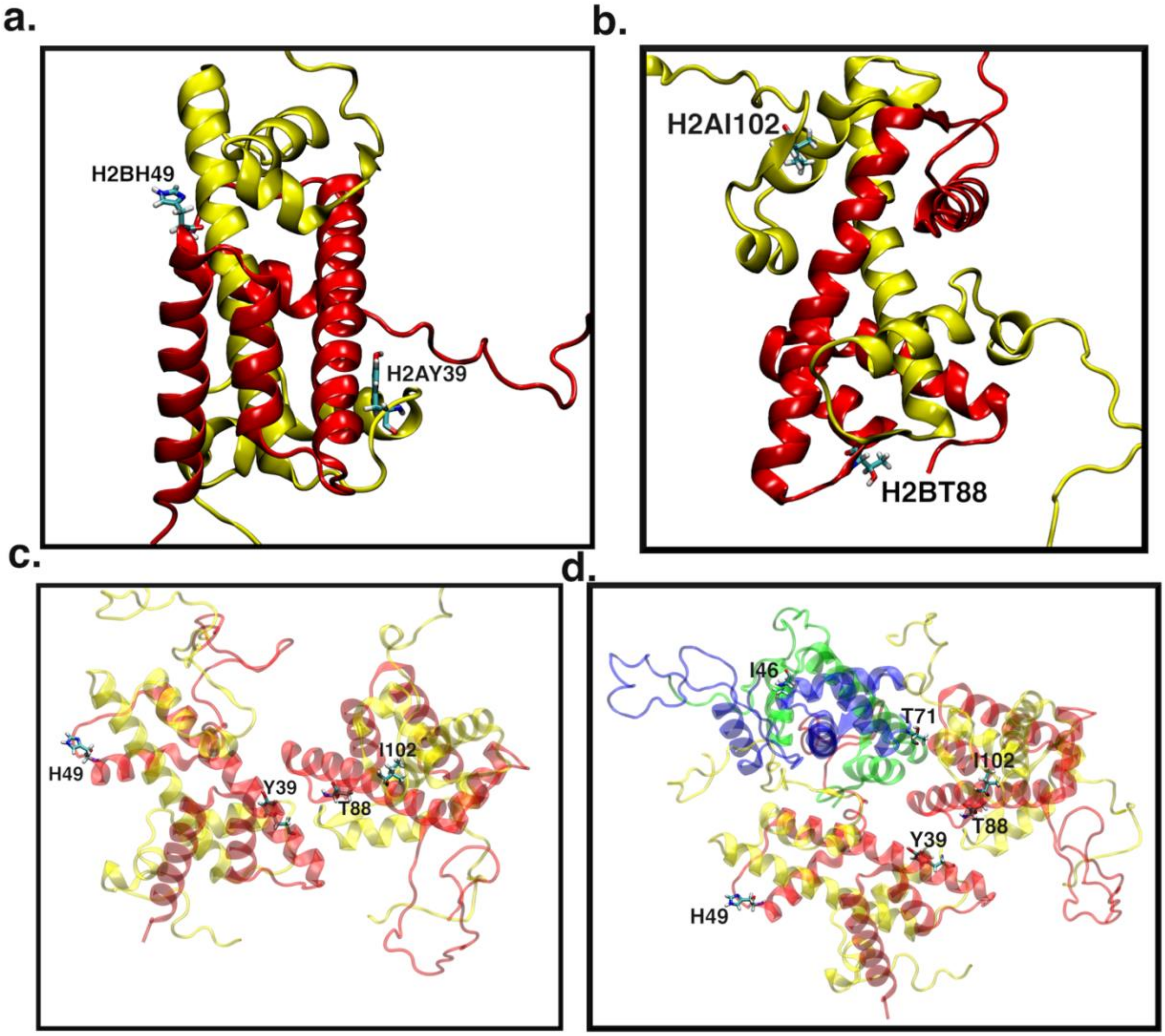


**Suppl. Figure 3**: Principal eigenvector showing kinetically connected domains in the nucleosome reveal major contributors to domain communication at (a) The first copy H2A-H2B dimer (b) The second copy H2A-H2B dimer (c) H2A-H2B dimers interface (d) The H2A-H2B dimers tetramer (H3-H4) interface. H2BH49, H2BT88, H2AY39, H2AI102, H4T71 and H4I46 were important for dimer-dimer and dimer-tetramer communication in both the 1KX5 and 3LZ0 systems.

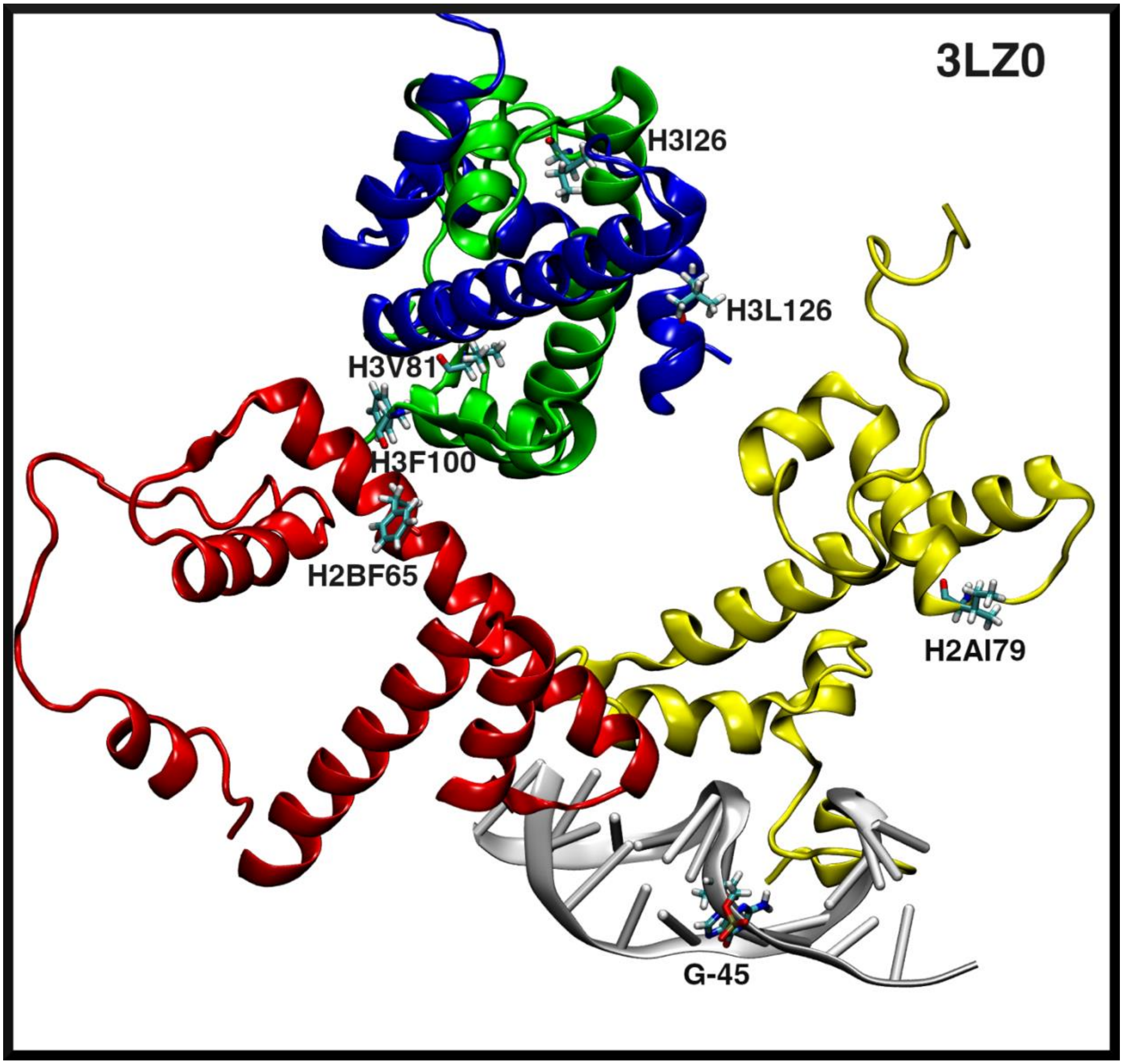


**Suppl. Figure 4**: Kinetic connected domains in the 3LZ0 systems among H2A-α3 helix (yellow), H2B-α2 helix (red), the SHL-4.5 region (grey). Tetramer-dimer kinetic connection was also seen at the α3 helix and C-terminus of H3 (blue) and loop 2 and α1 helix of H4 (green).

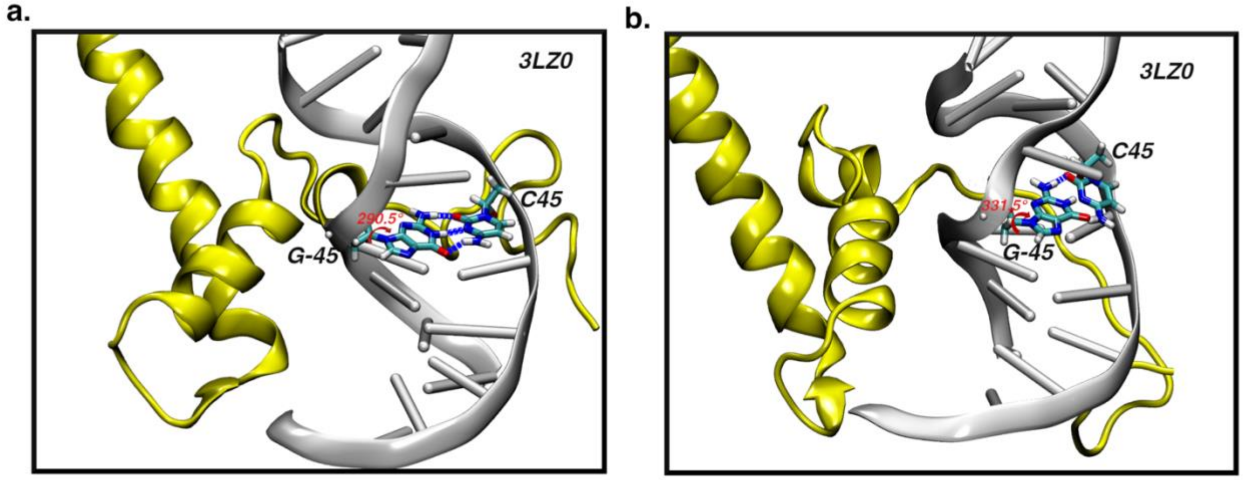


**Suppl. Figure 5**: Interaction between H2A.2 (yellow) and the SHL-4.5 region (grey) in the 3LZ0 systems strains the sugar-base dihedral angle in G-45 ($3^{I}$-$5^{I}$ strand) from (a) 290.5° to (b) 331.5°. This strain affected the Watson-Crick base pairing between G-45 ($3^{I}$-$5^{I}$ strand) and C45 ($5^{I}$-$3^{I}$ strand).

**Table S1: Inter-residue Conditional Activity in the 1KX5 System**

| Residue_X | Mut_Site_X | PTM_Site_X | Residue_Y | Mut_Site_Y | PTM_Site_Y | A[X][Y] |
|---|---|---|---|---|---|---|
| H2A-ARG11 | YES | NO | H2A-GLU121 | YES | NO | 5.1390582 |
| H2A-ARG11 | YES | NO | H2B-GLU76 | YES | NO | 5.0827703 |
| H2A-ARG11 | YES | NO | H3-ARG26 | YES | YES | 4.8070789 |
| H2A-ARG11 | YES | NO | H3-GLU105 | YES | NO | 5.12733925 |
| H2A-ARG11 | YES | NO | H3-LYS27 | YES | YES | 5.61304305 |
| H2A-ARG11 | YES | NO | H3-LYS36 | YES | YES | 5.14852455 |
| H2A-ARG17 | YES | NO | H2B-GLU76 | YES | NO | 4.9941249 |
| H2A-ARG17 | YES | NO | H3-LYS27 | YES | YES | 4.4862282 |
| H2A-ASN38 | YES | NO | H2A-GLU121 | YES | NO | 4.3000319 |
| H2A-ASN38 | YES | NO | H2B-GLU76 | YES | NO | 4.48938495 |
| H2A-ASN38 | YES | NO | H3-LYS27 | YES | YES | 4.65159407 |
| H2A-GLU56 | YES | NO | H2A-GLU121 | YES | NO | 4.6178368 |
| H2A-GLU56 | YES | NO | H2B-GLU76 | YES | NO | 4.6489866 |
| H2A-GLU56 | YES | NO | H3-GLU105 | YES | NO | 4.7053004 |
| H2A-GLU56 | YES | NO | H3-LYS27 | YES | YES | 5.50078415 |
| H2A-GLU56 | YES | NO | H3-LYS36 | YES | YES | 4.5238165 |
| H2A-HIE31 | YES | NO | H2A-GLU121 | YES | NO | 5.38488705 |
| H2A-HIE31 | YES | NO | H2B-GLU76 | YES | NO | 6.21782425 |
| H2A-HIE31 | YES | NO | H3-ARG26 | YES | YES | 4.93370575 |
| H2A-HIE31 | YES | NO | H3-GLU105 | YES | NO | 4.88700103 |
| H2A-HIE31 | YES | NO | H3-GLU73 | YES | NO | 4.162218 |
| H2A-HIE31 | YES | NO | H3-LYS27 | YES | YES | 6.25109653 |
| H2A-HIE31 | YES | NO | H3-LYS36 | YES | YES | 5.12125988 |
| H2A-LYS75 | YES | NO | H2A-GLU121 | YES | NO | 4.9014803 |
| H2A-LYS75 | YES | NO | H2B-GLU76 | YES | NO | 4.94398455 |
| H2A-LYS75 | YES | NO | H3-GLU105 | YES | NO | 4.8806861 |
| H2A-LYS75 | YES | NO | H3-LYS27 | YES | YES | 5.04049998 |
| H2A-LYS75 | YES | NO | H3-LYS36 | YES | YES | 5.0151929 |
| H2B-ASP68 | YES | NO | H2A-GLU121 | YES | NO | 4.64501627 |
| H2B-ASP68 | YES | NO | H2B-GLU76 | YES | NO | 4.81621008 |
| H2B-ASP68 | YES | NO | H3-ARG26 | YES | YES | 4.1161991 |
| H2B-ASP68 | YES | NO | H3-LYS27 | YES | YES | 4.79239777 |
| H2B-ASP68 | YES | NO | H3-LYS36 | YES | YES | 4.622401 |
| H2B-PHE70 | YES | NO | H2A-GLU121 | YES | NO | 5.00311883 |
| H2B-PHE70 | YES | NO | H2B-GLU76 | YES | NO | 5.5414222 |
| H2B-PHE70 | YES | NO | H3-ARG26 | YES | YES | 4.8077568 |

| | | | | | | |
|---|---|---|---|---|---|---|
| H2B-PHE70 | YES | NO | H3-GLU105 | YES | NO | 4.99378957 |
| H2B-PHE70 | YES | NO | H3-LYS27 | YES | YES | 5.55266098 |
| H2B-PHE70 | YES | NO | H3-LYS36 | YES | YES | 4.9052639 |
| H3-ARG131 | YES | NO | H2A-GLU121 | YES | NO | 4.87284128 |
| H3-ARG131 | YES | NO | H2B-GLU76 | YES | NO | 5.76448445 |
| H3-ARG131 | YES | NO | H3-ARG26 | YES | YES | 4.4752468 |
| H3-ARG131 | YES | NO | H3-GLU105 | YES | NO | 4.62611047 |
| H3-ARG131 | YES | NO | H3-LYS27 | YES | YES | 5.1521137 |
| H3-ARG131 | YES | NO | H3-LYS36 | YES | YES | 4.4697732 |
| H3-ARG42 | NO | YES | H2A-GLU121 | YES | NO | 5.2093579 |
| H3-ARG42 | NO | YES | H2B-GLU113 | YES | NO | 5.17035525 |
| H3-ARG42 | NO | YES | H2B-GLU76 | YES | NO | 6.9241259 |
| H3-ARG42 | NO | YES | H3-ARG26 | YES | YES | 5.61925893 |
| H3-ARG42 | NO | YES | H3-GLU105 | YES | NO | 5.15230463 |
| H3-ARG42 | NO | YES | H3-GLU73 | YES | NO | 7.4158872 |
| H3-ARG42 | NO | YES | H3-LYS27 | YES | YES | 6.4898276 |
| H3-ARG42 | NO | YES | H3-LYS36 | YES | YES | 6.55136993 |
| H3-GLU73 | YES | NO | H3-GLU73 | YES | NO | 4.3509735 |
| H4-ARG45 | YES | NO | H2A-GLU121 | YES | NO | 4.3908732 |
| H4-ARG45 | YES | NO | H2B-GLU76 | YES | NO | 4.67316055 |
| H4-ARG45 | YES | NO | H3-GLU105 | YES | NO | 4.1407869 |
| H4-ARG45 | YES | NO | H3-LYS27 | YES | YES | 5.0886667 |
| H4-ARG45 | YES | NO | H3-LYS36 | YES | YES | 4.2920779 |
| H4-ARG92 | YES | YES | H2A-GLU121 | YES | NO | 4.38530153 |
| H4-ARG92 | YES | YES | H2B-GLU76 | YES | NO | 4.67851345 |
| H4-ARG92 | YES | YES | H3-GLU105 | YES | NO | 4.3717737 |
| H4-ARG92 | YES | YES | H3-LYS27 | YES | YES | 4.42378157 |
| H4-GLU53 | YES | NO | H2A-GLU121 | YES | NO | 4.3851455 |
| H4-GLU53 | YES | NO | H2B-GLU76 | YES | NO | 5.2406152 |
| H4-GLU53 | YES | NO | H3-GLU105 | YES | NO | 4.1478119 |
| H4-GLU53 | YES | NO | H3-LYS27 | YES | YES | 4.7082665 |
| H4-GLU53 | YES | NO | H3-LYS36 | YES | YES | 4.2580818 |
| H4-LEU49 | YES | NO | H3-LYS27 | YES | YES | 4.2494276 |
| H4-SER47 | NO | YES | H2B-GLU76 | YES | NO | 4.104216 |
| H4-SER47 | NO | YES | H3-LYS27 | YES | YES | 4.23842363 |

**Table S2: Predicted Kinetically Connected domain with Residues contributing to Principal Eigenvector and Dynamical Memory in the NCP Systems**

| Histone | Domain | Contributors to eigenvector | High Dynamical memory |
|---|---|---|---|
| **H3** | αN helix | Tyr 54 | — |
| **H3** | αN–α1 loop/ α1 helix | Leu 61; Gln 76 | Arg 63 |
| **H3** | α2 helix | Phe 104 | — |
| **H3** | α3 helix | Leu 126; Ile 130; Arg 129; Arg 134 | — |
| **H4** | α1 helix | Ile 26; Ile 29; Thr 30; Arg 39 | Asp 24, Arg 35, Arg 36, Arg 40 |
| **H4** | Loop 1 | Ile 46; | Lys 44 |
| **H4** | α2 helix | Val 70; Thr 71; Tyr 72 | Glu 53, Arg 55 |
| **H4** | Loop 2 | Val 81 | — |
| **H4** | α3 helix | Phe 100 | Arg 92 |
| **H2A** | Loop 1 | Tyr 39 | — |
| **H2A** | α3 helix | Ile 78; Gln 84 | — |
| **H2A** | C-terminal tail/ docking domain | Glu 92; Ile 102 | — |
| **H2B** | Loop 1 | His 49; Ile 54 | — |
| **H2B** | α2 helix | Phe 65; Phe 70 | Phe 70 |
| **H2B** | Loop 2 | Tyr 83; Thr 88 | — |
| **H2B** | α3 helix | Ile 89; Leu 100 | — |
| **H2B** | αC-terminal tail | Thr 115 | — |
